# Semi-automated seizure detection using interpretable machine learning models

**DOI:** 10.1101/2023.10.25.563903

**Authors:** Pantelis Antonoudiou, Trina Basu, Jamie Maguire

## Abstract

Numerous methods have been proposed for seizure detection automation, yet the tools to harness these methods and apply them in practice are limited. Here we compare four interpretable and widely-used machine learning models (decision tree, gaussian naïve bayes, passive aggressive classifier, stochastic gradient descent classifier) on an extensive electrographic seizure dataset collected from chronically epileptic mice. We find that the gaussian naïve bayes model achieved the highest precision and f1 score, while also detecting all seizures in our dataset and only requires a small amount of data to train the model and achieve good performance. We use this model to create an open-source python application SeizyML that couples model performance with manual curation allowing for efficient and accurate detection of electrographic seizures.

**Author Summary:** Seizure detection based on electrographic recordings is critical for epilepsy diagnosis and research. However, the current gold standard for seizure detection is manual curation, which is biased, costly, incredibly laborious, and requires extensive training and expertise, prohibiting advances in epilepsy diagnosis and research. Here we demonstrate that fast, simple, and interpretable machine learning (ML) models are sufficient to detect all seizures in an extensive dataset collected from a well-established mouse model of epilepsy. Importantly, we created an open-source python application, SeizyML, that integrates the most precise model tested here with human curation of the detected events. This semi-automated approach greatly enhances efficiency and precision of seizure detection while also being transparent. We believe that the adoption of semi-automated and transparent technologies is indispensable for understanding ML model predictions, improving their reliability, and fostering trust between ML models and neurodiagnostic professionals.

## Introduction

Epilepsy is a devastating neurological disorder characterized by recurrent seizures. Detection of seizures by evaluation of the electroencephalogram (EEG) is critical for diagnosis and Epilepsy research (Flink et al., 2002). However, manual detection, which is the current standard of practice, is laborious, error prone, requires extensive training and expertise, and results in large variability (Diachenko et al., 2022). Artificial intelligence (AI) and Machine learning (ML) techniques hold great promise in assisting clinical diagnosis and transforming research (Rajpurkar et al., 2022; Yu et al., 2018); as can be seen by the prompt FDA approval of AI tools (Benjamens et al., 2020).

A wealth of ML methods have been proposed for automating the task of seizure detection (Shoeibi et al., 2021; Siddiqui et al., 2020). Specifically, deep learning techniques automate feature extraction and perform remarkably well in seizure detection among other tasks (Cho & Jang, 2020; Rajpurkar et al., 2022; Shankar et al., 2022; Shoeibi et al., 2021; Yuan et al., 2019). However, deep learning techniques are not easily interpretable (Rudin, 2019). This is an important issue as many clinicians and scientists may be reluctant to employ these methodologies due to lack of trust (Pinto et al., 2022; Rudin, 2019); as the characterization of parameters that constitute a seizure is not clear. Furthermore, deep learning models often require a large amount of training data to reach a good performance and can be prohibitive to train due to expensive computational resources (Rajpurkar et al., 2022; Yu et al., 2018). Finally, these existing approaches make it difficult to extract the electrographic features which may be informative regarding seizure generation and progression.

Here we have employed widely-used and interpretable machine learning models from the scikit-learn python library (Pedregosa et al., 2011) and tested their performance on detecting seizures from a well-established mouse model of epilepsy. We found that these interpretable models with combined simple feature extraction were sufficient to detect all seizures in our dataset. To make these pipelines accessible to the scientific community we created an open-source python application (SeizyML) to couple high model sensitivity with manual verification of the detected seizures. This semi-automated approach significantly reduces time required for seizure detection while providing high sensitivity and accuracy.

## Methods

### EEG/LFP Dataset

The data used here for training, validation, and testing of the models were obtained from recordings collected in our previous study (Basu et al., 2022). Briefly, adult mice were injected with kainic acid in the ventral hippocampus (vHPC) and were then implanted with a stainless-steel wire in the vHPC and a stainless-steel screw fixed above frontal cortex (FC). Data were sampled at 4000 samples per second. The data were split into training (11 mice, 4224 hours), and testing (15 mice, 5511 hours) datasets. Subsequently data were divided in 5 second windows and downsampled to 100 Hz with an antialiasing filter, as this has been shown to achieve excellent performance (Cho & Jang, 2020; Jang & Cho, 2019). Data were also high pass filtered at 2 Hz to remove baseline drift and extreme outliers (>25 standard deviations) and were replaced with the median of each 5 second window. Then 9 features were extracted for each channel (autocorrelation, line length, root mean square, mean absolute deviation, variance, standard deviation, power (2-40Hz), energy, envelope amplitude) and 3 features were extracted for cross channel metrics (cross-correlation, covariance, absolute covariance) (Table 1) that resulted in a total of 21 features (9 for each channel + 3 cross channel). The features were converted into z-scores to equalize their contribution to the machine learning models.

**Table 1.**
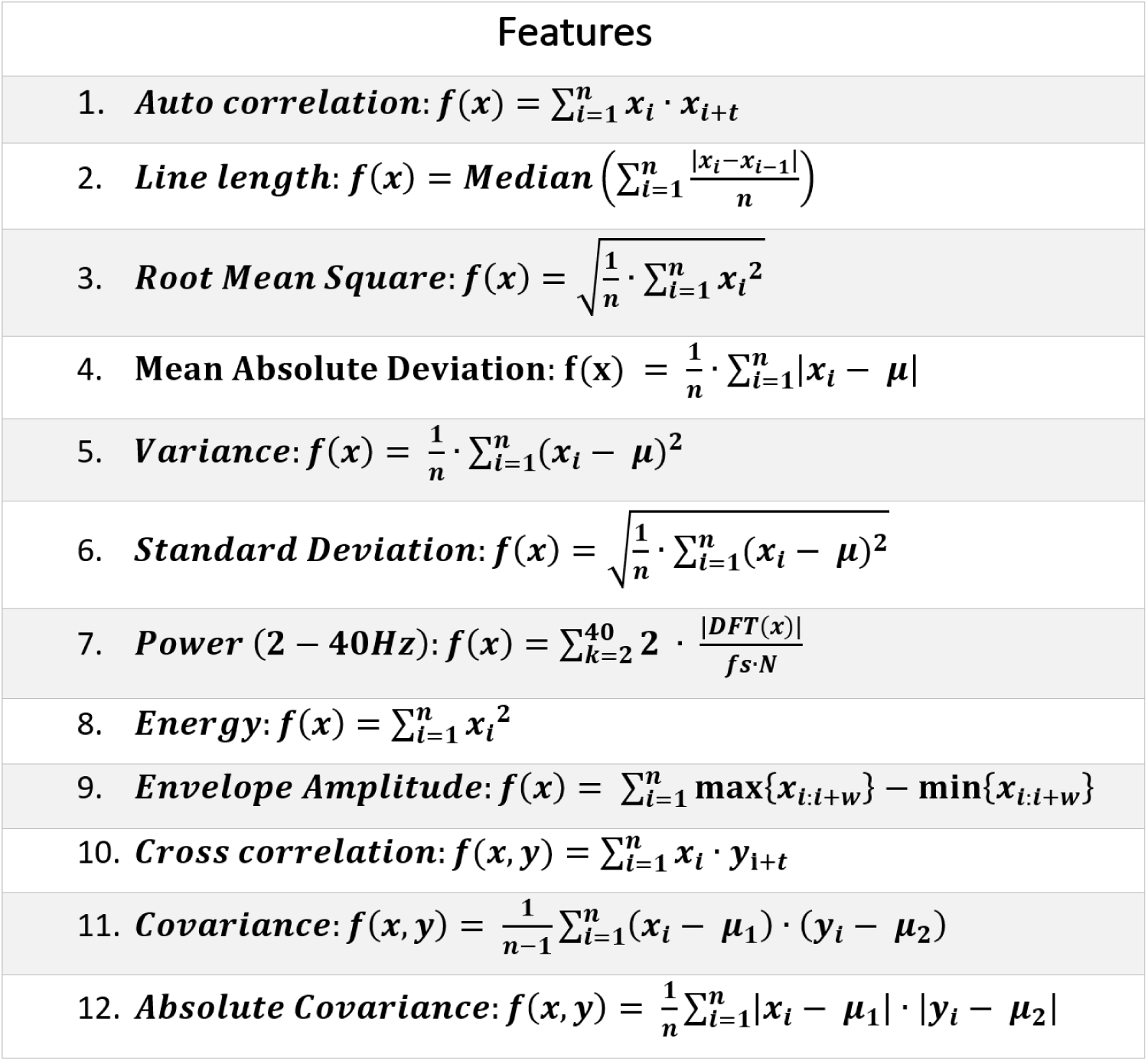
Features. Where **n** is the length of the signal (**n**=500 samples, 5 seconds), **i** is the index, **DFT** is the Discrete Fourier Transform (from SciPy’s *fft*), **fs** is the sampling rate (**fs**=100 Hz), **μ** is the mean, **t** is the time lag (**t**=0), and **w** is the window (**w**=30 samples, 0.3 seconds).

### Feature Selection

In order to remove redundant information and select the most meaningful features we first removed highly correlated features with Pearson coefficient *r* >0.99, which resulted in 12 features that constituted the first feature-set (Table 2, Column 1). We then selected the 4 and 8 best ranking features from ANOVA comparison with target variable (Table 2, Column 2&3) and from mutual information (Table 2, Column 4&5), resulting in a total of 5 feature-sets (Table 2).

**Table 2:** Feature selection. Columns represent feature-sets. Cells that contain 1 in a blue background indicate that the feature was selected. Columns **1-5)** Highly correlated features were removed. Columns **2-3)** Best four and eight ranked features selected by ANOVA. Columns **4-5)** four and eight ranked features selected by mutual information.

|  | Feature Set | 1 | 2 | 3 | 4 | 5 |
| --- | --- | --- | --- | --- | --- | --- |
| vHPC | Auto-correlation vHPC | 1 | 0 | 1 | 0 | 1 |
|  | Line Length vHPC | 1 | 1 | 1 | 1 | 1 |
|  | Root Mean Square vHPC | 1 | 0 | 0 | 0 | 1 |
|  | Mean Abs. Dev, vHPC | 1 | 0 | 1 | 1 | 1 |
|  | Variance vHPC | 0 | 0 | 0 | 0 | 0 |
|  | Standard Deviation vHPC | 0 | 0 | 0 | 0 | 0 |
|  | Power (2-40 Hz) vHPC | 0 | 0 | 0 | 0 | 0 |
|  | Energy vHPC | 0 | 0 | 0 | 0 | 0 |
|  | Envelope Amplitude vHPC | 1 | 0 | 0 | 1 | 1 |
| FC | Auto-correlation FC | 1 | 1 | 1 | 0 | 0 |
|  | Line Length FC | 1 | 1 | 1 | 1 | 1 |
|  | Root Mean Square FC | 1 | 0 | 0 | 0 | 0 |
|  | Mean Abs. Dev. FC | 1 | 0 | 1 | 0 | 1 |
|  | Variance FC | 0 | 0 | 0 | 0 | 0 |
|  | Standard Deviation FC | 0 | 0 | 0 | 0 | 0 |
|  | Power (2-40 HZ) FC | 0 | 0 | 0 | 0 | 0 |
|  | Energy FC | 0 | 0 | 0 | 0 | 0 |
|  | Envelope Amplitude FC | 1 | 0 | 1 | 0 | 1 |
| vHPC-FC | Cross-correlation | 1 | 0 | 0 | 0 | 0 |
|  | Covariance | 0 | 0 | 0 | 0 | 0 |
|  | Absolute Covariance | 1 | 1 | 1 | 0 | 0 |

### Feature Contributions

Feature contributions to model predictions were obtained as follows:

1. For the DT model, we retrieved feature importance from the trained DT models (Sklearn *DecisionTreeClassifier)*, where feature importance was calculated as the Gini importance.
2. For the SGD model, we calculated feature weight as the absolute weights for each feature from the trained SGD model (Sklearn *SGDClassifier*).
3. For the GNB model, we calculated feature separation score based on extracted parameters of each class from the trained GNB model (Sklearn *GaussianNB*):

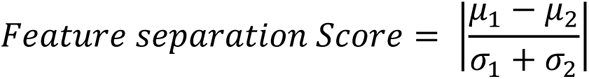

Where *μ* is the mean and *σ* is the standard deviation for each class.

*\*Feature contributions were normalized to have a sum of 1 for each model metric*.

### Model Selection and Training

We selected four models for seizure detection, gaussian naïve bayes (GNB), decision tree (DT), stochastic gradient descent classifier (SGD) and passive aggressive classifier (PAC) from the scikit-learn (Sklearn) toolbox (Pedregosa et al., 2011). We selected models based on interpretability and their ability to train on large datasets. The SGD model was used with log/hinge loss to implement logistic regression/support vector machine models, respectively. This was done as logistic regression and support vector machine models could not train effectively on large datasets. The PAC model was selected as another implementation of a support vector machine model (hinge loss) with a different optimization algorithm (Crammer et al., 2006). The k-nearest neighbors model was not used as it was too slow and therefore not useful in practice during model predictions. The models were then tuned by performing grid-search using the balanced accuracy as the fit metric to optimize their hyperparameters. Hyperparameter search was performed using 4-fold cross validation (75% training subset, 25% validation subset) of the training dataset (11 mice, 4224 hours). Sklearn’s *StratifiedKfold* method was used to ensure that the class distributions were similar in training and validation datasets. The tuning was performed for each model, for every feature-set, and the resulting best hyperparameters were selected (Table 3). Each model/feature-set combination from Table 3 was then trained 5 times using *StratifiedKfold* (5 folds, 80% training subset) and evaluated on the testing dataset (15 mice, 5511 hours).

**Table 3:**
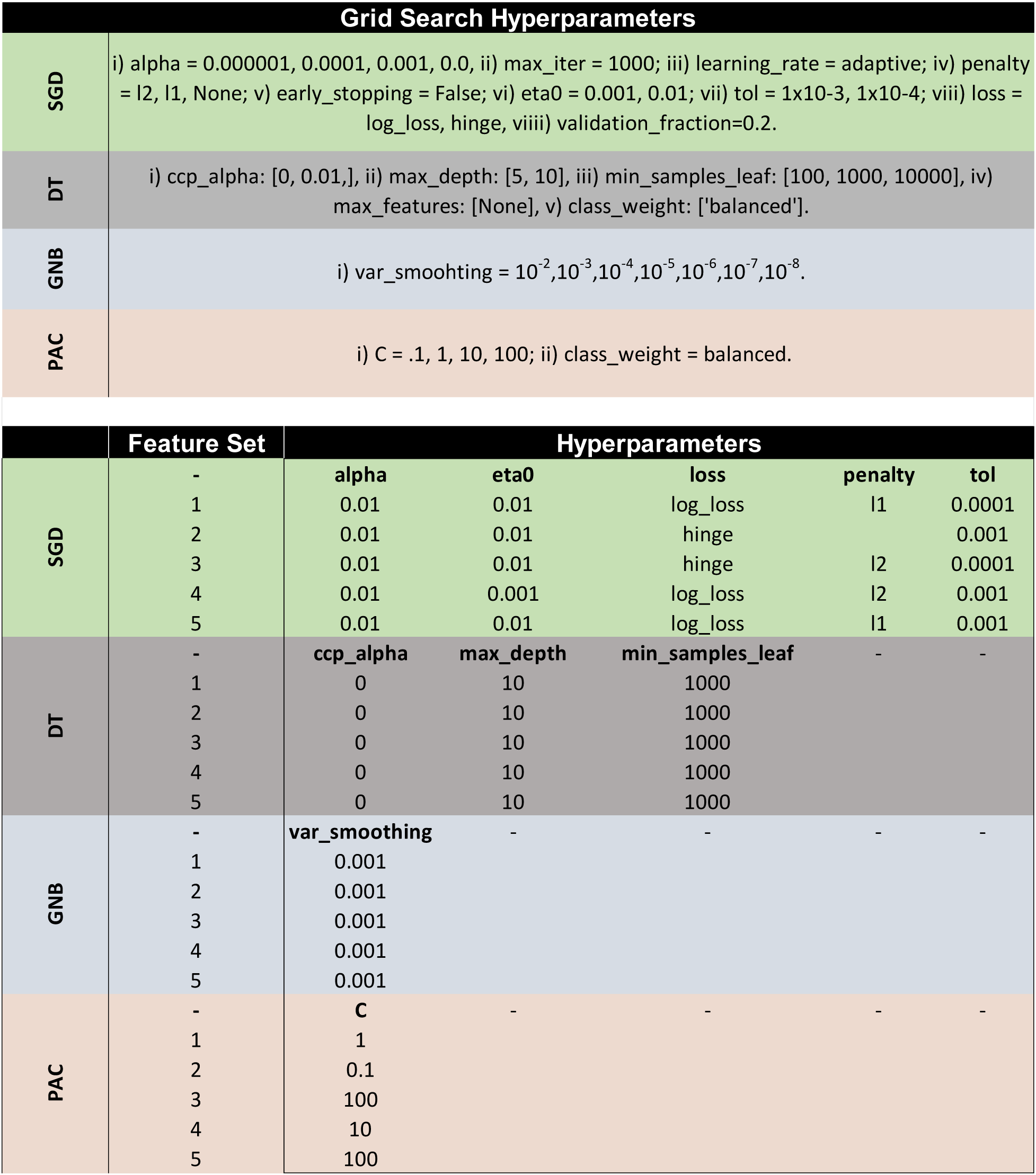
Hyperparameter optimization. **Top)** Hyperparameter space used to perform grid search. **Bottom)** Hyperparameters that resulted in best balanced accuracy scores were selected for each model and feature-set. Note that only hyperparameters with more than one value from the **top** are displayed on the **bottom** table.

| Grid Search Hyperparameters |  |
| --- | --- |
| SGD | i) alpha = 0.000001, 0.0001, 0.001, 0.0, ii) max_iter = 1000; iii) learning_rate = adaptive; iv) penalty = l2, l1, None; v) early_stopping = False; vi) eta0 = 0.001, 0.01; vii) tol = 1x10-3, 1x10-4; viii) loss = log_loss, hinge, ix) validation_fraction=0.2. |
| DT | i) ccp_alpha: [0, 0.01], ii) max_depth: [5, 10], iii) min_samples_leaf: [100, 1000, 10000], iv) max_features: [None], v) class_weight: ['balanced']. |
| GNB | i) var_smoothing = 10<sup>-2</sup>, 10<sup>-3</sup>, 10<sup>-4</sup>, 10<sup>-5</sup>, 10<sup>-6</sup>, 10<sup>-7</sup>, 10<sup>-8</sup>. |
| PAC | i) C = .1, 1, 10, 100; ii) class_weight = balanced. |

**Table 3: Hyperparameter optimization.**
|  | Feature Set | Hyperparameters |  |  |  |  |
| --- | --- | --- | --- | --- | --- | --- |
| SGD | - | alpha | eta0 | loss | penalty | tol |
|  | 1 | 0.01 | 0.01 | log_loss | l1 | 0.0001 |
|  | 2 | 0.01 | 0.01 | hinge |  | 0.001 |
|  | 3 | 0.01 | 0.01 | hinge | l2 | 0.0001 |
|  | 4 | 0.01 | 0.001 | log_loss | l2 | 0.001 |
|  | 5 | 0.01 | 0.01 | log_loss | l1 | 0.001 |
| DT | - | ccp_alpha | max_depth | min_samples_leaf | - | - |
|  | 1 | 0 | 10 | 1000 |  |  |
|  | 2 | 0 | 10 | 1000 |  |  |
|  | 3 | 0 | 10 | 1000 |  |  |
|  | 4 | 0 | 10 | 1000 |  |  |
|  | 5 | 0 | 10 | 1000 |  |  |
| GNB | - | var_smoothing | - | - | - | - |
|  | 1 | 0.001 |  |  |  |  |
|  | 2 | 0.001 |  |  |  |  |
|  | 3 | 0.001 |  |  |  |  |
|  | 4 | 0.001 |  |  |  |  |
|  | 5 | 0.001 |  |  |  |  |
| PAC | - | C | - | - | - | - |
|  | 1 | 1 |  |  |  |  |
|  | 2 | 0.1 |  |  |  |  |
|  | 3 | 100 |  |  |  |  |
|  | 4 | 10 |  |  |  |  |
|  | 5 | 100 |  |  |  |  |

### Model Metrics

1. True Positive (TP): Segments that were correctly predicted as seizures.
2. True Negative (TN): Segments that were correctly predicted as non-seizures.
3. False Positive (FP): Segments that were predicted as seizures but are non-seizures.
4. False Negative (FN): Segments that were predicted as non-seizures but are seizures.

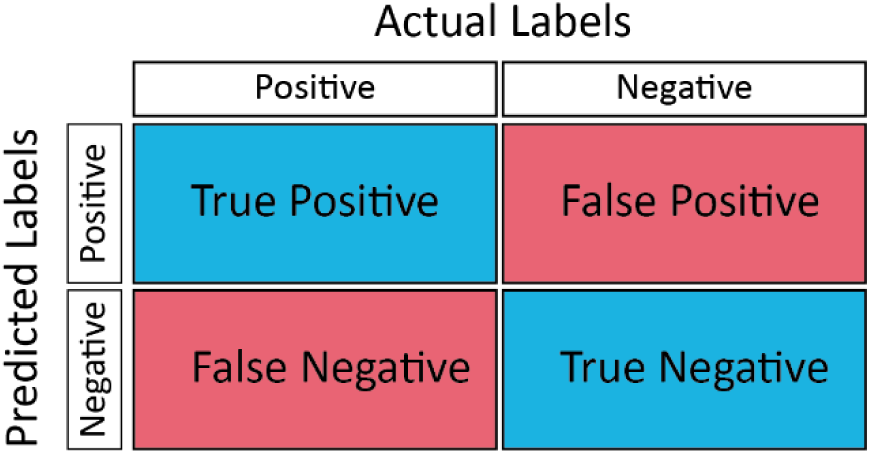
5. 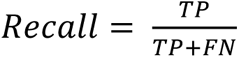: Proportion of identified positives.
6. 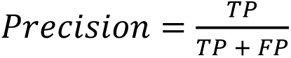: Proportion of predicted positives that were actually positive.
7. 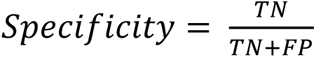: Proportion of identified negatives.
8. 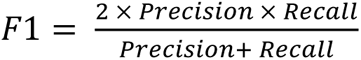
9. 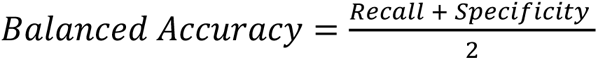
10. 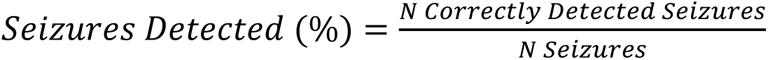
11. 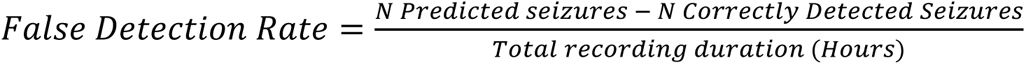

*\*Detected segments were defined as seizures if there were at least two consecutive 5 segments*.

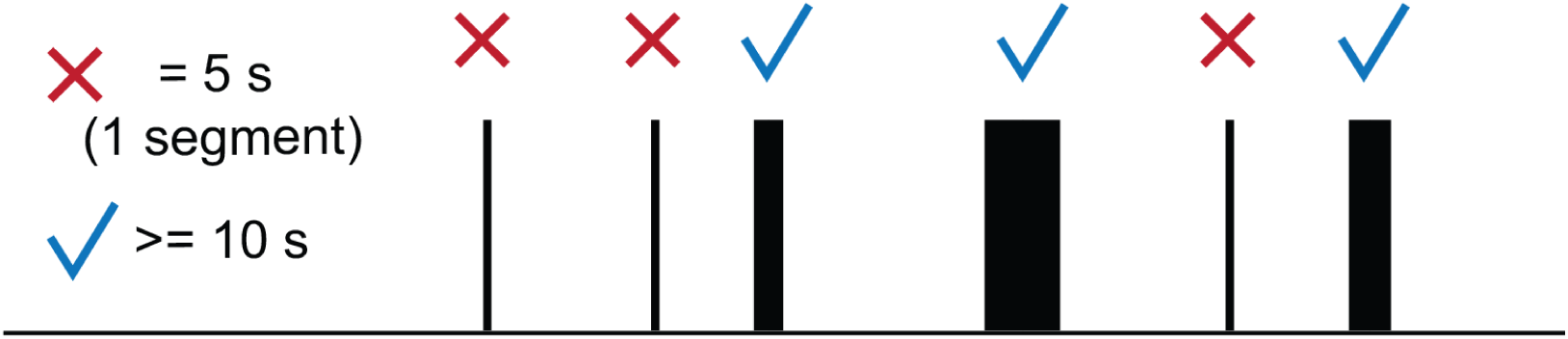

### Statistical Tests

To compare the metric across models a one-way ANOVA was used with pairwise Tukey’s HSD multiple comparisons using the python toolbox *statsmodels*. All bar and line plots represent the mean and error-bars, or shaded regions represent the SEM. A p-value <0.05 was considered statistically significant.

### Code

The open-source seizure detection app, seizyML, can be found on GitHub at https://github.com/neurosimata/seizy_ml. All other code will be made available upon reasonable request.

## Results

An examination of even a few seizures in a well-established and reproducible model allows us to appreciate the variability and diversity of these events (Figure 1a). This variability becomes even more apparent when looking at extracted features (See methods – feature selection) across multiple seizures (Figure 1b). Even though the extracted features robustly increase during seizure events, their variability during seizure events is greatly enhanced when compared to the periods before and after seizure events (Figure 1b). Due to the inherent variability in seizure events, we trained machine learning models using these extracted features and examined their seizure detection performance.

**Figure 1:**
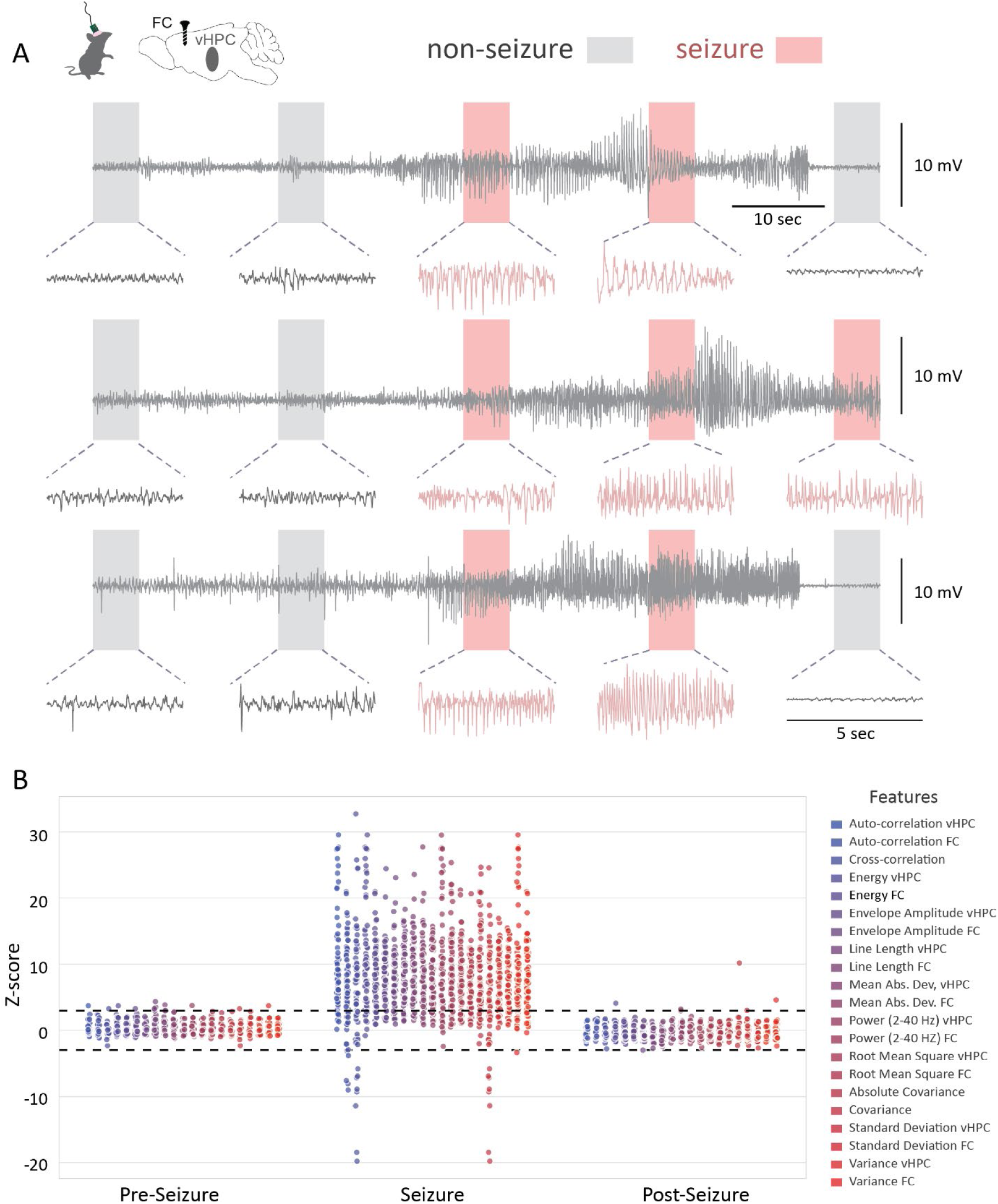
Example seizures and extracted features from the training dataset. **A)** Example seizure and non-seizure traces from ventral hippocampus (vHPC) LFP. Each row represents one seizure that was obtained from a different recording (n = 3 recordings, 2 animals), **B)** Dot plot of all extracted features before, during and after seizure from training dataset. Dotted horizontal lines indicate Z-scores of +/- 3.

To achieve this, we split the data into training (11 mice, 4224 hours), and testing (15 mice, 5511 hours) datasets. We then selected 5 feature-sets (Table 1) by removing redundant features and quantifying their relevance (Table2, See methods – feature selection). After feature selection, we chose four models (Figure 2A) for seizure detection: decision tree (DT), gaussian naïve bayes (GNB), passive aggressive classifier (PAC), and stochastic gradient descent classifier (SGD) based on model interpretability and ability to efficiently train on our dataset (See methods – model selection). Each model was tuned (Table 3, hyperparameter selection) and then trained 5 times for each feature-set (Table 2) to account for model variability and to obtain a better estimate of their performance.

**Figure 2:**
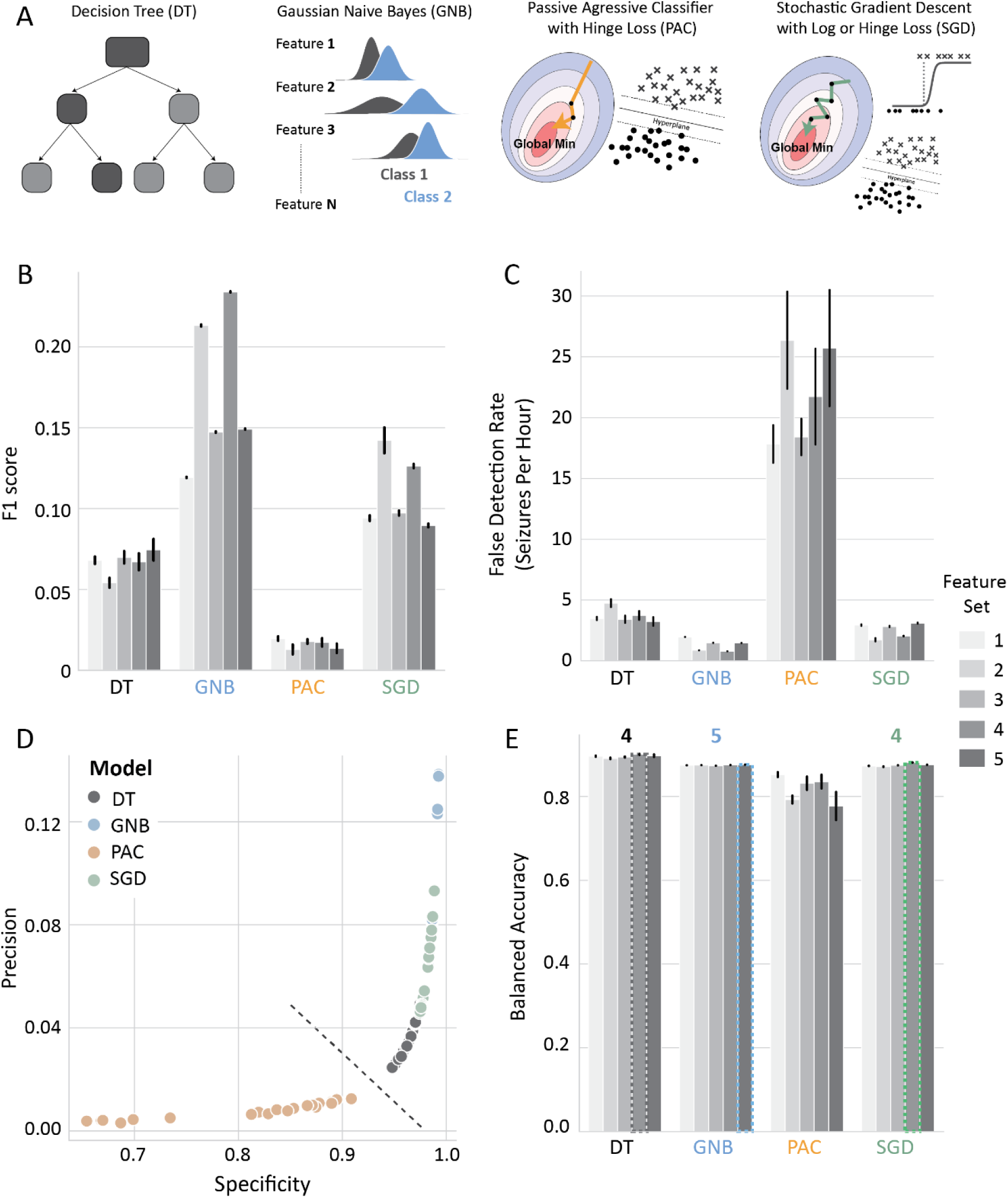
Comparison of ML models across feature-sets. **A)** Schematic of ML models compared, **B)** F1 score**, C)** False detection rate: Number of incorrectly detected seizures per hour, **D)** Scatterplot of precision vs specificity; dotted line indicates the separation of PAC with the rest of the models, **E)** Selection of feature-set from Table 2 based on balanced accuracy; Numbers on top of bars indicate the selected feature-set per model.

We first performed a comparison of the 4 models across all feature combinations (Figure 2A-B). We observed that the PAC model had substantially lower F1 score (Figure 2B) – a combined measure of model precision and sensitivity (See methods – model metrics) and high false detection rate compared to the other 3 models independent of the feature combination (Figure 2C). All the PAC trained models had lower precision and specificity across all models (Figure 2D). Therefore, the PAC model was not considered for further analysis. For each of the three remaining models, the feature combination that resulted in the highest balanced accuracy was selected for further examination. Specifically, feature combinations 4, 5, 4 were chosen for DT, GNB, and SGD, respectively (Figure 2E).

Next, we compared the DT, GNB, and SGD models across several metrics (Figure 3A). Overall, the DT model had the lowest number of false negatives – incorrectly classified seizure segments as non-seizures (Figure 3B; DT = 147.87 ± 12.40 x10^3^, GNB = 52.87 ± 0.16 x10^3^, SGD = 65.69 ± 0.81 x10^3^) resulting in the highest recall (Figure 3C; DT = 0.84 ± 0.002, GNB = 0.77 ± 0.000, SGD = 0.78 ± 0.001) among the three models. However, it also had the highest number of false positives – incorrectly classified non-seizure segments as seizures (Figure 3D; DT = 1.00 ± 0.013 x10^3^, GNB = 1.46 ± 0.001 x10^3^, SGD = 1.38 ± 0.003 x10^3^) resulting in the lowest precision (Figure 3E; DT = 0.03 ± 0.003, GNB = 0.08 ± 0.000, SGD = 0.07 ± 0.001). These results indicate that the DT model was the most sensitive and the least precise among the three models. On the other hand, the GNB model was the most precise (Figure 3E) and had the highest F1 score (Figure 3F; DT = 0.07 ± 0.005, GNB = 0.15 ± 0.000, SGD = 0.13 ± 0.001). The SGD model had an intermediate performance overall with a lower F1 score than the GNB model (Figure 3F). Even though the DT model has the highest recall, it had a similar if not slightly worse performance at seizure detection (See methods – model metrics) than GNB and SGD models which detected all seizures in the test dataset (Figure 3G; DT = 99.80 ± 0.033 %, GNB = 100.00 ± 0.000 %, 100.00 ± 0.000). These results demonstrate that simple and interpretable machine learning models can be very efficient for seizure detection but vary in their reliability and prediction accuracy.

**Figure 3:**
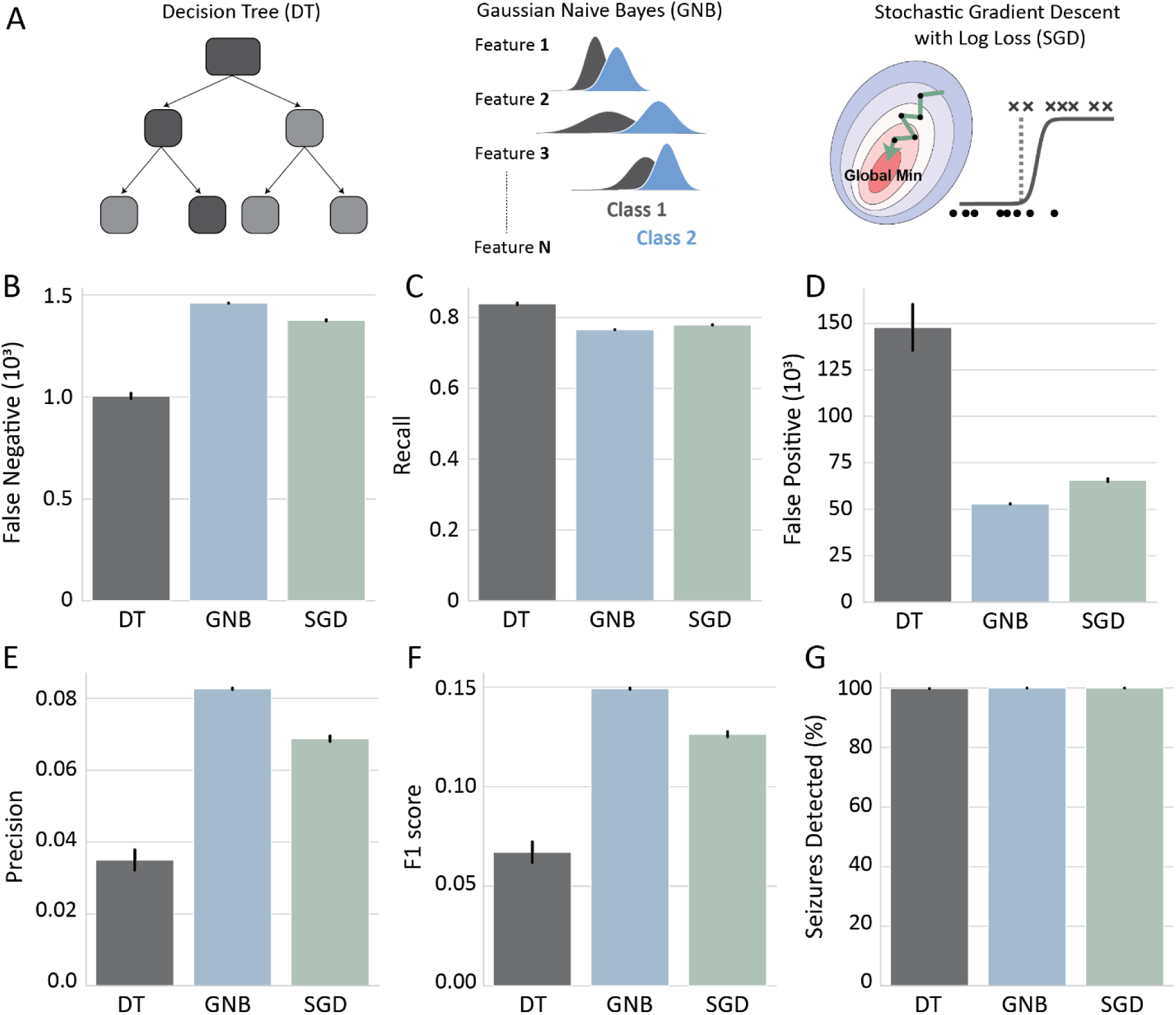
Comparison of selected ML models. **A)** Schematic of selected machine learning models based on balanced accuracy (Figure 2E). **(B-G)** Comparison of select models across metrics **B)** False Negatives, **C)** Recall, **D)** False Positives, **E)** Precision, **F)** F1 score, **G)** Percentage of seizures detected from the testing dataset.

**Supplemental Table 1:**
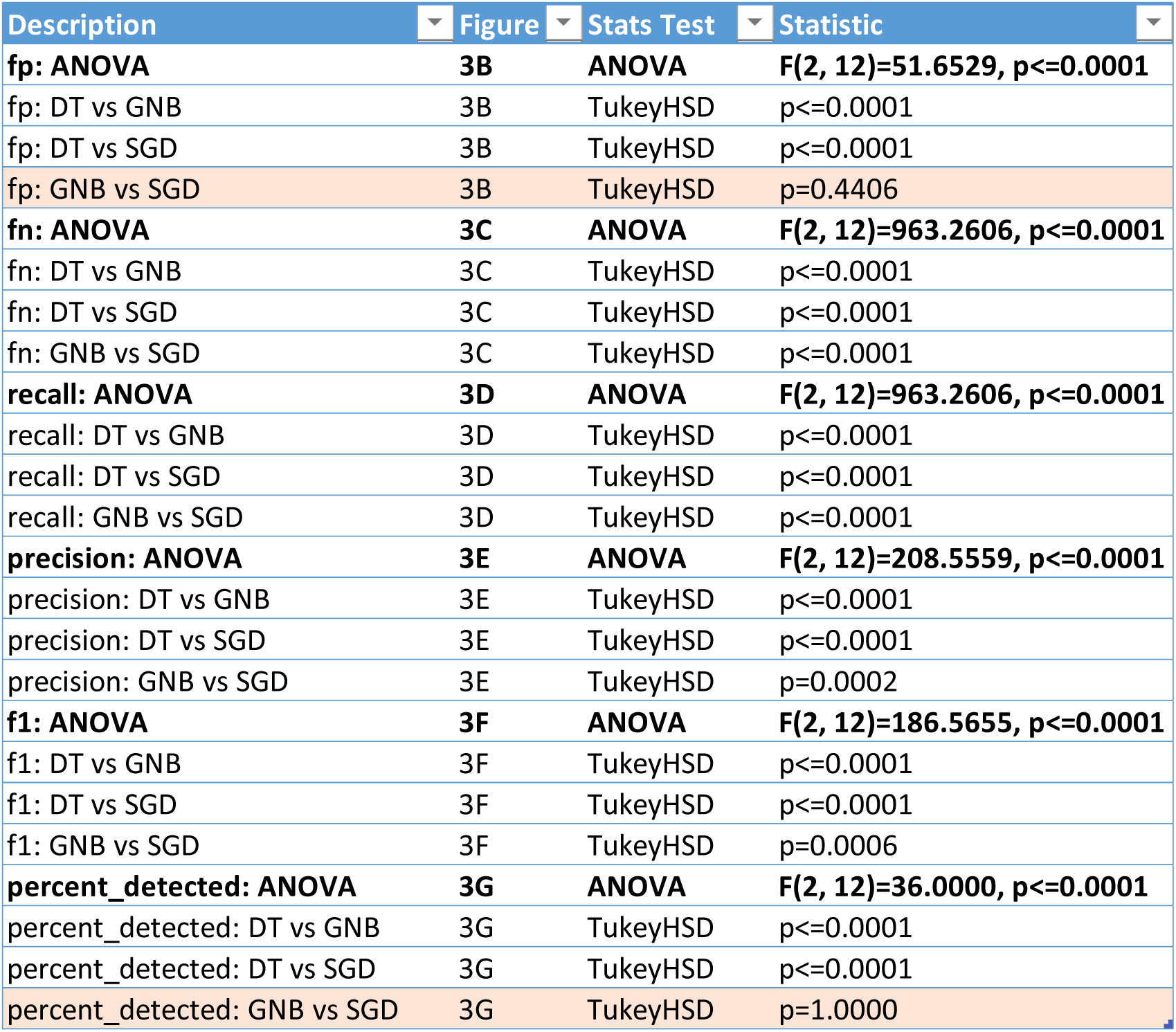
Statistical comparison of 3 models across metrics. Non-significant p-values (p > 0.05) are highlighted in orange.

Even though the DT model had significantly higher recall than the GNB model, the number of seizures detected was not superior (DT: 99.80 ± 0.03%, GNB: 100.00 ± 0.00%). Given that the recall of all three models was lower than the proportion of seizures detected, we investigated how the predicted seizure bins across time compared between the three models and ground truth data. When comparing the predicted seizure bins to ground truth data, we observed that the models detected the center of the seizure with higher accuracy than seizure boundaries (Figure 4A-C). This is not surprising given that the features that were used to train these models do not increase as robustly at the designated seizure boundaries (Supplemental Figure 1). This observation could explain why the proportion of segments predicted correctly as seizures is lower than the proportion of detected seizures. Interestingly, it seems that the increased recall of the DT model arises from high detection of seizure offset segments. However, the DT model dramatically misclassifies seizure offset as false positives (Figure 4A, D), which likely accounts for its decreased precision. This was not specific to the selected feature-set chosen to train the DT model or the depth of the tree, as all DT models tested here overestimated seizure offset predictions (Supplemental Figure 2).

**Figure 4:**
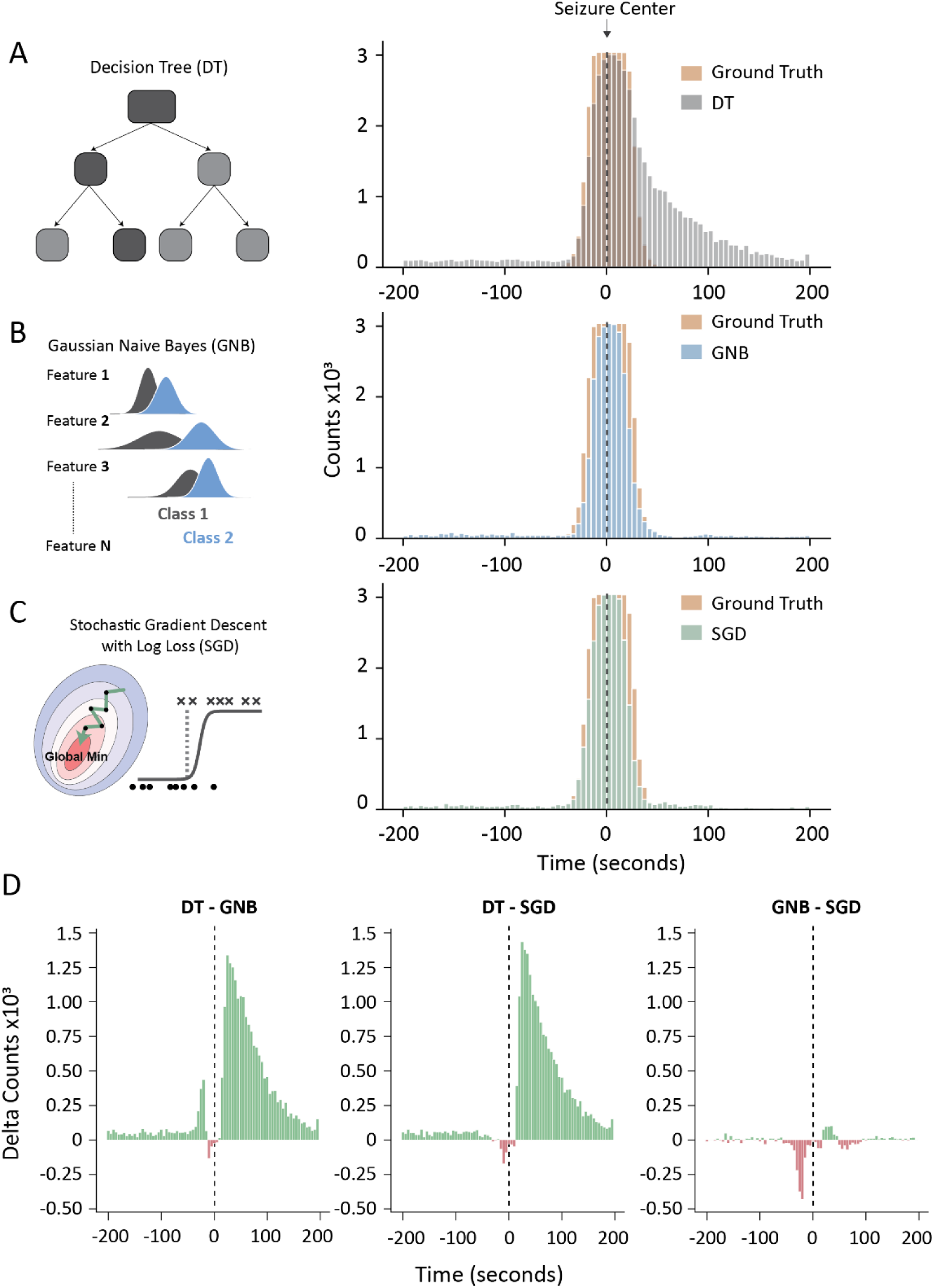
Seizure prediction across time vs ground truth data. **(A-C)** Number of ground truth vs predicted seizure across time bins. **A)** DT, **B)** GNB, **C)** SGD. **D)** Difference between model predictions across the 3 pairs. Dotted line represents the seizure center.

**Supplemental Figure 1.**
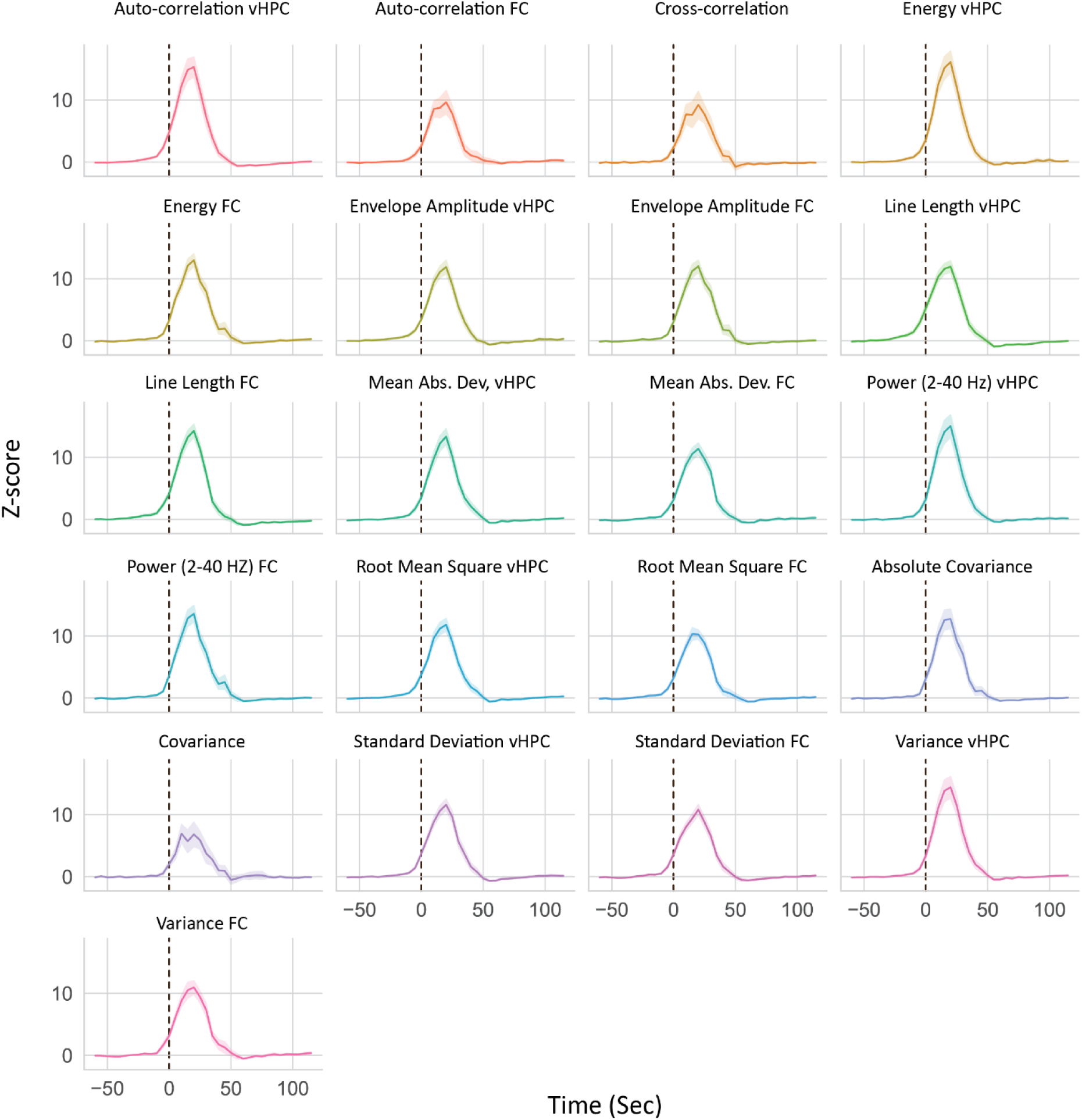
Feature values change seizure onset. Graphs show how normalized feature values increase during time-locked seizure onset across all seizures in the training dataset. Seizure onset is marked by black dotted line.

**Supplemental Figure 2:**
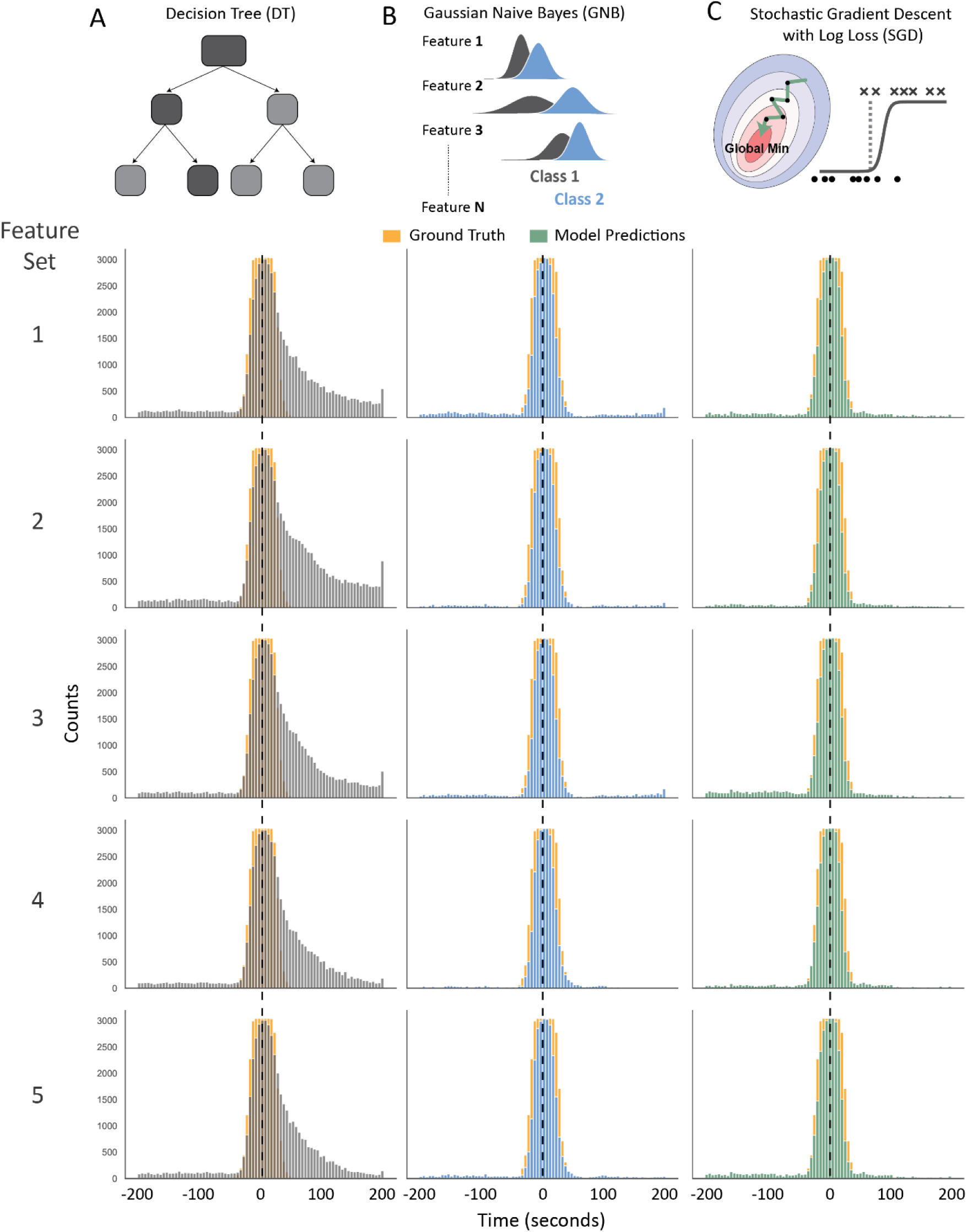
Seizure prediction across time vs ground truth data. **(A-C)** Number of ground truth vs predicted seizure across features-sets. **A)** DT, **B)** GNB, **C)** SGD. Dotted line represents the seizure center.

Manual inspection of EEG datasets to create training labels is costly and laborious. To examine the dataset size required to achieve good model performance and seizure detection, we trained models on increasing size of data portions (Figure 5). The GNB model detected all seizure events from just 1% of the training data even though its performance based on F1 score and balanced accuracy, seems to stabilize at 10% of the training data (Figure 5A-B). The SGD model detected 99.74 % of all seizures at 1% of training data and detected all seizures at 2.5% of the training data whereas its F1 score seems to have stabilized around 10% of the training data, although consistently below the GNB model (Figure 5A-B). The DT model detected 99.93% of all seizures at 1% of training data and detected all seizures at 2.5% of the training data, but its detection dropped to 99.84% at 100% of training data size (Figure 5A). This likely resulted from a DT model optimization to increase precision and reduce false positives (Figure 5C-D), resulting in higher number of false negatives (Figure 5E). In addition, the F1 score of the DT model kept improving as the training data size increased up to the full dataset. However, the F1 score, and false detection rate (Figure 5D, F), were lower than the GNB and SGD models across training data sizes. As observed before, the DT model had a much lower number of false negatives in comparison to the SGD and GNB models even though it had a reduced seizure detection overall (Figure 5A, E). Therefore, the GNB and SGD model perform well even with smaller training data sizes and quickly achieve a stable performance. On the other hand, the DT model requires a large amount of data to improve its precision at the cost of reduced seizure detection.

**Figure 5:**
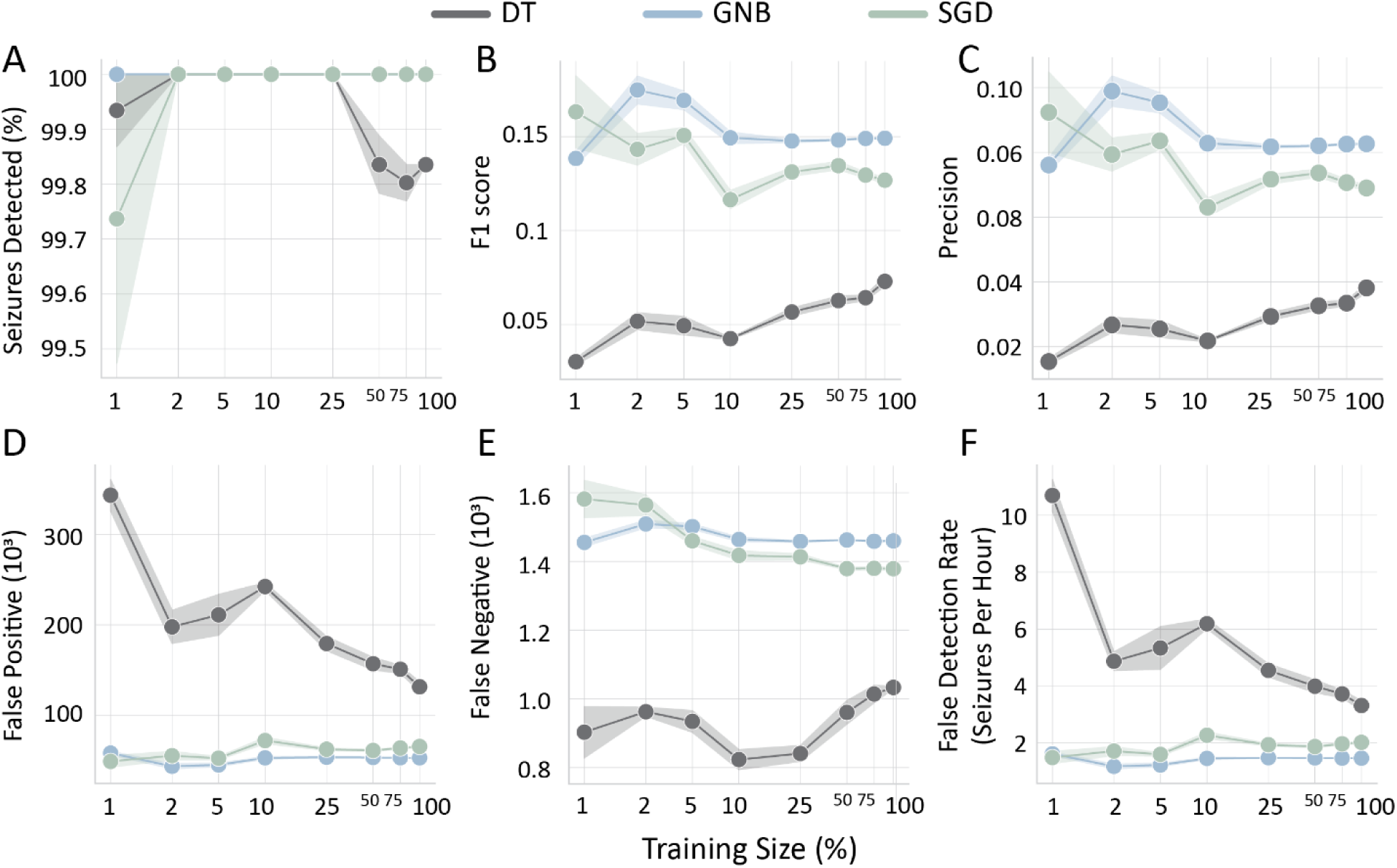
Model performance as a function of training data size. Performance of ML models across key metrics for increasing portions of the training dataset. **A)** Percent seizures detected, **B)** F1 score, **C)** Precision, **D)** False Positives, **E)** False Negatives, **F)** False detection rate: Number of incorrectly detected seizures per hour.

To further understand how these models classified EEG segments we extracted metrics which quantified the influence of each feature on model predictions (See methods: feature contributions; DT: feature importance, SGD: feature weight, GNB: feature separation score). This analysis revealed that in the DT model the *line length of the vHPC* was by far the most important feature with a value of 0.80. The second most important feature was the *envelope amplitude of the vHPC* with only a value of 0.14, while the two other features had negligible importance, each scoring less than 0.05 (Figure 6A). In contrast the SGD model had a more balanced weight across features, with the *line length of the vHPC* also having the highest weight score of 0.40. The *envelope amplitude of the vHPC* was a close second, with a feature weight of 0.36, while the other two features had a combined score of 0.24 (Figure 6B). Lastly, the GNB model does not have an in-built metric for feature importance thus we calculated a feature separation score based on the distribution of each feature from the trained GNB models (See methods – feature contributions). Our analysis indicated that most features had comparable scores, although the *line length of vHPC* had a marginally higher score (Figure 6C). Thus, the GNB model appears to have a more balanced feature contribution for its predictions. Overall, this analysis reveals that the *line length of vHPC* feature is a key contributor to seizure detection in this dataset, whereas feature contributions varied across models. Intriguingly, the models with more balanced feature contributions also had a superior performance.

**Figure 6:**
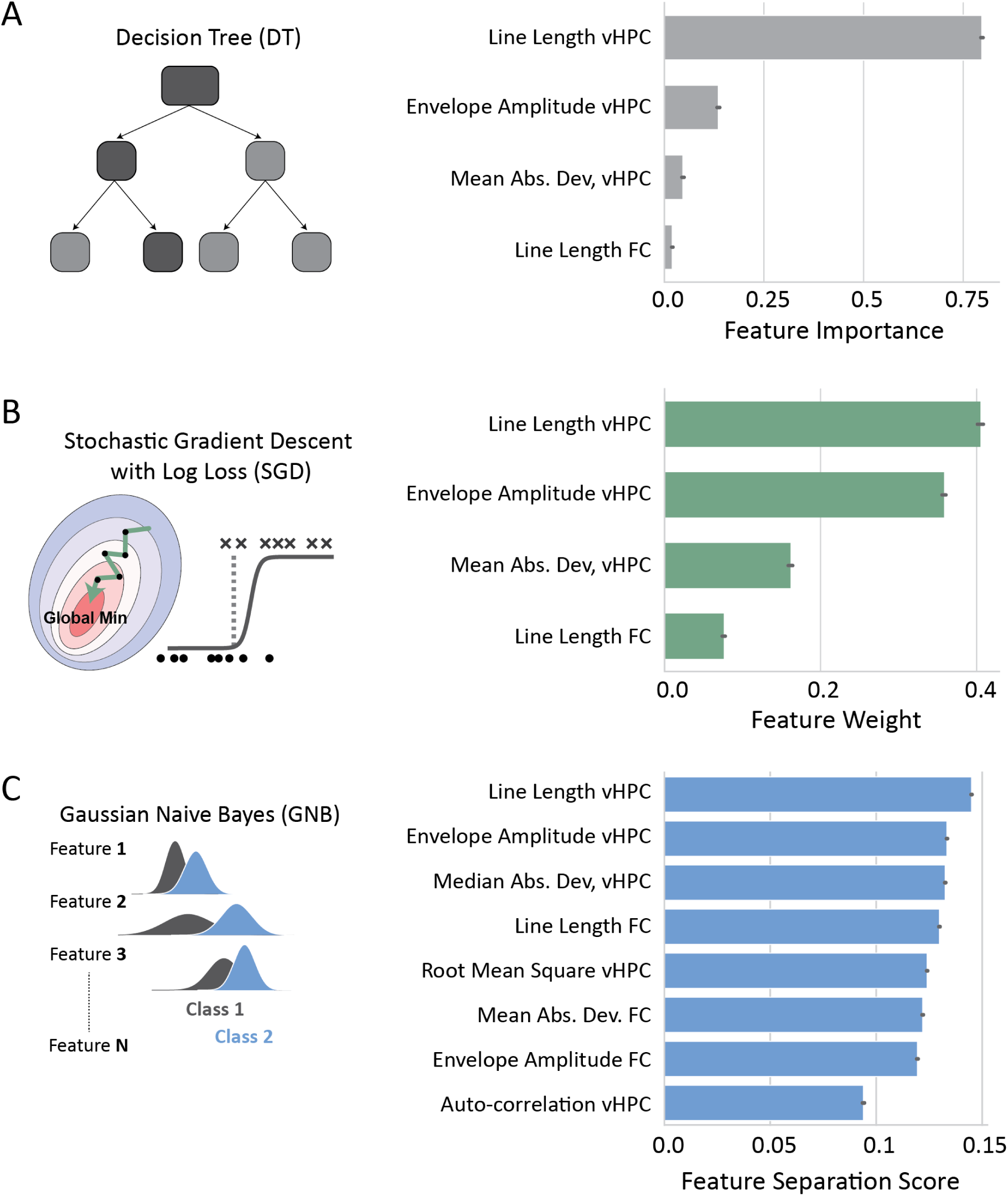
Feature importance for classification. **A)** DT: Feature importance, **B)** SGD: Feature weight, **C)** GNB: Feature separation score. Each feature-set has a cumulative sum of 1.

### A semi-automated application for seizure detection

Here we observed that interpretable ML models with simple feature extraction were very efficient at detecting seizures in our mice that were injected with kainic acid in the ventral hippocampus (Basu et al., 2022). To couple the high model sensitivity with enhanced accuracy we created an open-source application for semi-automated seizure detection, SeizyML, that combines model predictions with manual curation of the detected seizure events. The outline of the pipeline is illustrated in Figure 7. Before the app can be used, the raw LFP/EEG data should be downsampled (100 Hz, 5 second windows) and must be converted from their native format (depending on recording apparatus) to HDF5. A small training dataset needs to also be prepared to train the gaussian naïve bayes model. Then using the command line interface of SeizyML, the data are preprocessed, features are extracted, and model predictions are generated. Following that a simple GUI allows the user to accept or reject the detected seizures. Lastly, seizure properties can be extracted from the detected seizures using the seizyML CLI. Importantly, the app can be easily extended to use any machine learning (ML) model, channel number and features. Although, care should be taken since some ML models cannot perform well on large datasets especially on large number of features (including decision tree models used here).

**Figure 7:**
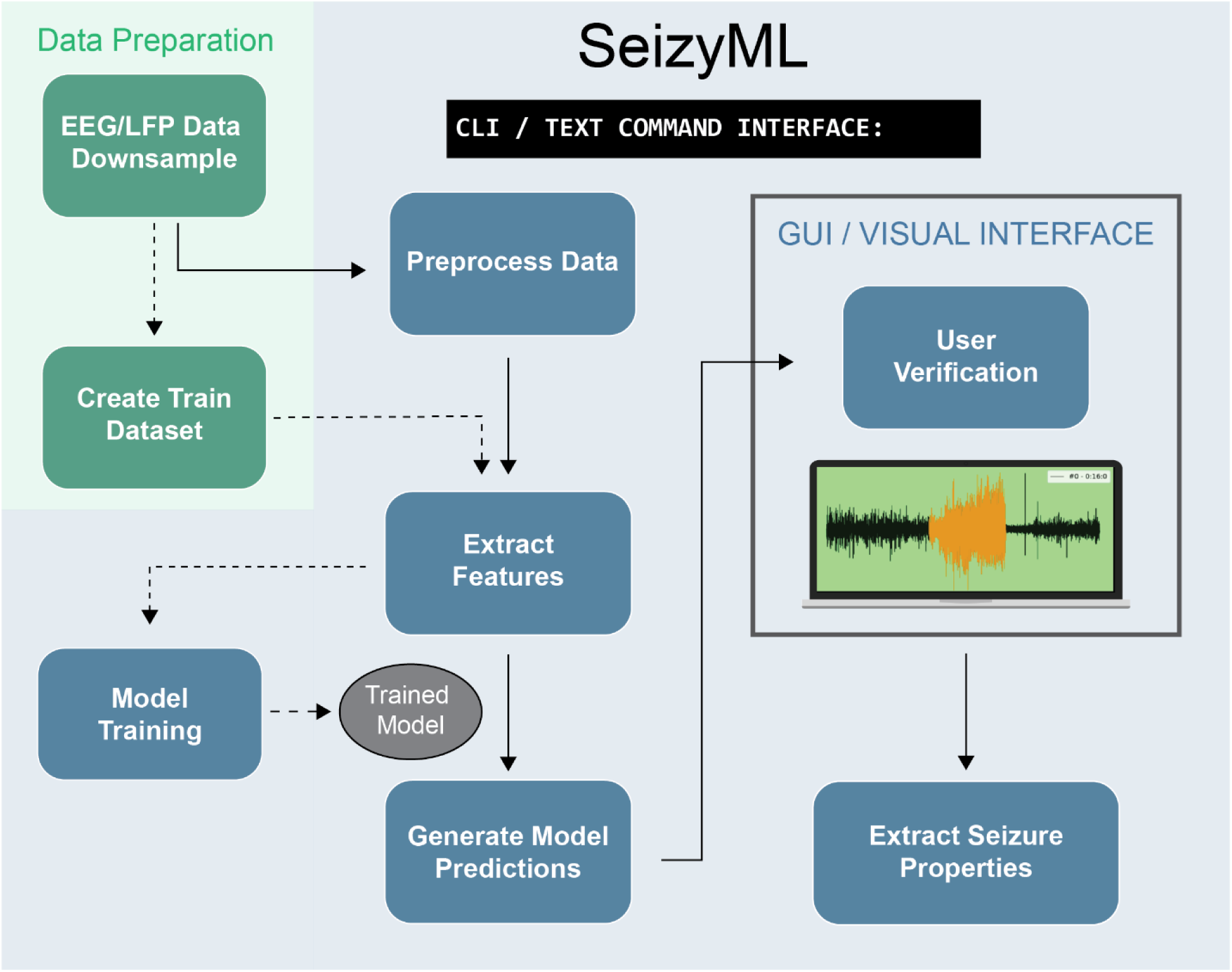
SeizyML pipeline. Green area denotes data preparation steps performed from the user before the data are ready for processing by SeizyML. Blue area highlights the main processing steps of the SeizyML application.

## Discussion

Here we have compared the performance of four linear models for automated seizure detection: gaussian naïve bayes (GNB), decision tree (DT), stochastic gradient descent classifier (SGD) and passive aggressive classifier (PAC). Even though most models detected all seizures in our test dataset (608 seizures), we found that the GNB model had the best F1 score, had simple tuning (only one hyperparameter), needed only a small amount of training data to reach good performance and it is highly interpretable. In agreement with previous findings (Logesparan et al., 2012), we found that the best predictor for seizure detection was the line length of the electrographic activity (vHPC). This was revealed by 1) the feature ranks of the ANOVA/mutual information test from the training dataset along with 2) feature importance of the trained models. We have further created an open-source python application, SeizyML, for semi-automated seizure detection using the GNB model. We believe that a semi-automated approach that combines fast and sensitive machine learning models with seizure verification from human operators is a crucial first step in the automation of seizure detection. Allowing operators to review the detected seizures and verify them can enhance their trust of the algorithm and will help to understand how to further improve ML algorithms.

Our model comparisons revealed that the PAC model had the worst performance across multiple metrics independent of the feature-set that it was trained on (Figure 2). It’s possible that the suboptimal performance of the PAC model arises from the aggressive optimization algorithm that misses the global minimum. This is supported by the fact that the SGD classifier with a different optimization algorithm, but the same loss function (Table 3) outperformed the PAC model (Figure 2). The DT model had the highest recall and lowest precision when compared to DT and SGD models (Figure 3). However, the high recall of the DT model did not translate to better seizure prediction (Figure 2). Both the high recall of the model and low precision seems to arise from misclassifications of the seizure termination (Figure 4) and the poor performance of the model could be attributed to it heavily relying on one feature (Figure 6). The SGD model had comparable performance to the GNB model although it had a lower F1 score (Figure 3). Additionally, the SGD model has many hyperparameters and took the longest to tune using grid search, making it more complicated and time-consuming to train. Nevertheless, it will be interesting to see a direct comparison of the performance of SGD and GNB models on other extended seizure datasets from well-established but diverse models of epilepsy.

## Limitations and Future Directions

This study aimed at finding simple and interpretable ML models for offline seizure detection. Our method using 5 second windows is not able to detect the seizure onset with high temporal precision. If detecting seizure onset/offset is of primary interest, our model could be combined with additional time-sensitive algorithms such as wavelet transform or Empirical mode decomposition approaches (Chen et al., 2017; Molnár et al., 2023). Although, these algorithms tend to be computationally expensive so it would be inefficient to be used in isolation. Furthermore, here we downsampled the LFP/EEG data to 100Hz and excluded high frequency components that could be important for seizure detection (Bragin et al., 2004; Molnár et al., 2023). In addition, we did not perform an exhaustive search of interpretable ML models due to time constraints. However, this approach was sufficient to detect all seizures in this dataset and has also resulted in fast and efficient ML pipeline.

One crucial consideration is that most automated seizure detection algorithms, including the ones here, are often restricted in classifications of non-seizure vs seizure periods. Yet, there is a broad range of seizures as categorized by clinical outcomes (such as focal, generalized tonic-clonic, etc.) (Fisher et al., 2017; McCallan et al., 2023). Detecting these different types of seizures is crucial in epilepsy as it can assist in enhanced diagnosis, more targeted therapies, and better control of seizures (McCallan et al., 2023). However, supervised approaches can only be as good as our definitions of these seizure types. And even though definitions of seizure types are improving, they are based exclusively on clinical symptoms (Fisher et al., 2017), and do not take into account the pathophysiology of neuronal networks and the mechanism of seizure generation. Therefore, improving seizure definitions based on the pathophysiology of neuronal networks is of paramount importance. We believe that creation of diverse seizure datasets (from multiple brain regions, recording systems, seizure types) with improved seizure definitions should be of utmost priority in order to improve epilepsy diagnosis and train better ML models for automation of seizure detection.

